# Predation Efficiency upon Clinical Isolates: *Bdellovibrio bacteriovorus* is Prey Specific and Origin Dependent

**DOI:** 10.1101/2021.07.13.452292

**Authors:** Claudia Saralegui, Cristina Herencias, Ana Halperin, Juan de Dios-Caballero, Blanca Pérez-Viso, Sergio Salgado-Briegas, Val F. Lanza, Rafael Cantón, Fernando Baquero, María Auxiliadora Prieto, Rosa del Campo

**Affiliations:** Department of Microbiology, Hospital Universitario Ramón y Cajal, Instituto Ramón y Cajal de Investigacion Sanitaria (IRYCIS), Madrid, Spain; Microbial Biotechnology Department, Centro Nacional de Biotecnología, CSIC, Madrid, Spain, C/ Darwin 3, 28049, Madrid Spain; Microbial and Plant Biotechnology Department, Biological Research Center-Margarita Salas, CSIC, Ramiro de Maeztu 9, 28040 Madrid, Spain

## Abstract

The use of predatory bacteria as live antibiotics has been proposed for managing bacterial infections, especially for those caused by antibiotic multiresistant isolates for which there are few therapeutic options. However, the current knowledge in this field is scarce, with most of the available data based on environmental isolates, with a significant lack of human clinical samples. In this study, we evaluated the predatory spectrum of the reference strain *Bdellovibrio bacteriovorus* 109J on 13 *Serratia marcescens* (5 of which were carbapenemase producers) and 78 *Pseudomonas aeruginosa* clinical isolates from respiratory (colonizing the lungs of patients with cystic fibrosis) or bacteremic infections, differentiated by phenotype (mucoid or not), antibiotic resistance phenotype (including multidrug-resistant isolates), and genetic lineage (frequent and rare sequence types). The source of the isolates was significantly associated with predation efficiency (100% for *S. marcescens,* 67% for *P. aeruginosa* from cystic fibrosis, and 25% for *P. aeruginosa* from bacteremia). In contrast, no correlation with colonial morphotype, genetic background, or antibiotic susceptibility was found. To evaluate the influence of the predator on the predation event, we employed a more aggressive *B. bacteriovorus* mutant 109J preying upon the same 48 bacteremic *P. aeruginosa* isolates. The mutant’s predation efficiency was higher than that of their wild-type counterpart (43% vs. 25%), pointing out that predation is specific to each prey-predator pair of isolates. Our results provide the most extensive study of clinical prey susceptibility published to date and show that the prey-predator interaction is influenced by the origin of the isolates rather than by their genetic background or their antibiotic susceptibility phenotype.

**IMPORTANCE:** The potential usefulness of predatory bacteria in controlling human pathogens, particularly those that are multiresistant to antibiotics, is enormous. Although this possibility has long been suggested, there are still no data on predation susceptibility in clinical strains, and the possible presence of autochthonous predators of the human microbiota has not been investigated. In this study, we employed a reference predator with an environmental origin to study predation phenomena in 3 well-characterized collections of human clinical isolates. Our results demonstrated that predation is a specific consequence of each prey-predator interaction, with the origin of the strains the most relevant factor. In contrast, the genetic background, morphotype, and antibiotic resistance did not appear to influence the predation phenomenon. We also highlight the involvement of a putative polyhydroxyalkanoate depolymerase protein of *B. bacteriovorus* in determining prey susceptibility. To our knowledge, this study is the largest performed with strains of clinical origin, discriminating between various genera and including strains with multiresistance to antibiotics.

## INTRODUCTION

The “golden age of antibiotics” in the mid-20th century was followed by the emergence of pathogens resistant to almost all available antibiotics, leading to the current global crisis of multidrug-resistant (MDR) bacteria (1, 2). To identify successful alternative antimicrobial therapies, bacterial pathogenicity needs to be understood as a multifactorial issue in which the surrounded microbiota, which includes natural competitors and predators, is also involved (3). In nature, predatory bacteria play an important role in maintaining population sizes by linking the production and removal of biomass in microbial communities, which in turn promotes the diversity of microorganisms and contributes to the global stabilization of the ecosystem (4, 5). The ecological role of predators could also be refined and exploited in the fight against clinical pathogens, given that the predators represent dynamic microorganisms that experience (as do their opponents) continuous physical, morphological, and metabolic adaptations, altering their behavior to counteract each other. This evolutionary reciprocity is the basis of coevolution, where adaptation by one player not only promotes change in its opponent, but the opponent’s adaptation likewise generates selection as an evolutionary response to the first player (6). Increasing our understanding of how the microbiota community ecology is balanced will contribute to the selection of biocontrol agents that target pathogenic bacteria for which antibiotics are not an alternative due to multiresistance (7–9). The successful development of predatory bacteria as “living antimicrobial agents” and a complete understanding of the predation mechanism depend on the characterization of their phenotypical predation preferences, mainly their prey range. A predator might have a wide repertoire of susceptible prey, but predation appears to be strongly strain-specific, fundamentally based on the composition of the prey cell envelope, and influenced by environmental conditions (10–12). However, addressing the specific factor driving prey preference and susceptibility is challenging and has so far remained elusive, particularly in non-environmental bacterial collections (13, 14).

The most studied bacterial predators are *Bdellovibrio* and like organisms (BALOs), which are small vibrioids, rod-shaped gram-negative aerobic bacteria, recently reclassified to the class of *Oligoflexia,* which belongs to the *Proteobacteria* phylum (15). Its known prey species spectrum includes genera of *Proteobacteria* phylum as *Pseudomonas, Escherichia, Acinetobacter, Aeromonas, Burkholderia, Citrobacter, Enterobacter,* and *Klebsiella* (11, 16–18), as well as antibiotic-resistant isolates (19). Although BALOs were first isolated from soil, they are ubiquitous in nature and can be found in aquatic and terrestrial environments, including hypersaline systems (20), biofilms (21), mammalian intestines (22–24), and the lungs of patients with cystic fibrosis (25). In addition to the genetic detection of BALOs, several authors have documented the *in vivo* phenomenon of predation in human microbial ecosystems (26–29).

Herein, we characterized the predation susceptibility and efficiency of the reference strain *B. bacteriovorus* 109J against human clinical *Serratia marcescens* and *Pseudomonas aeruginosa* isolates, of diverse origins, genetic backgrounds, and antibiotic-resistant phenotypes. We also explored the relevance of the predator role on predation using a more aggressive mutant, a previously described derivative of *B. bacteriovorus* 109J (30). We found a specific recognition of susceptible prey that could be related to its single deletion genotype in the bd2637 gene, coding for a putative polyhydroxyalkanoate (PHA) depolymerase enzyme.

## RESULTS

### Prey origin determines the predation susceptibility to *B. bacteriovorus* 109J

The use of predatory bacteria as biocontrol agents depends on their efficiency in eradicating bacterial populations and on which bacterial species are susceptible to predation, also known as the prey range. We measured the predation susceptibility of *S. marcescens,* CF-PA and BACT-PA clinical isolates by monitoring the decrease in OD_600_ of the predation co-cultures and by measuring the viable prey cell number.

All 13 *S. marcescens* isolates tested were preyed on by *B. bacteriovorus* 109J, whereas CF-PA isolates were significantly more susceptible to predation (20 out of 30, 67%) than BACT-PA (12 out of 48, 25%) (*p*<0.02) (Fig 1, S1, S2, and S3). No correlation between predation and *P. aeruginosa* genetic lineage was observed, with discrepancies in 6 of the 12 STs grouping more than one isolate [ST175 (1/3), ST253 (1/3), ST274 (1/2), ST532 (1/2), ST646 (2/3), and ST1017 (1/2)] (Table S1). There was also no correlation with the mucoid (4/7) or non-mucoid phenotype of CF isolates (16/23) (Fig S4) or with the antibiotic susceptibility phenotype (Fig 2 and Table S2).

**Fig 1.**
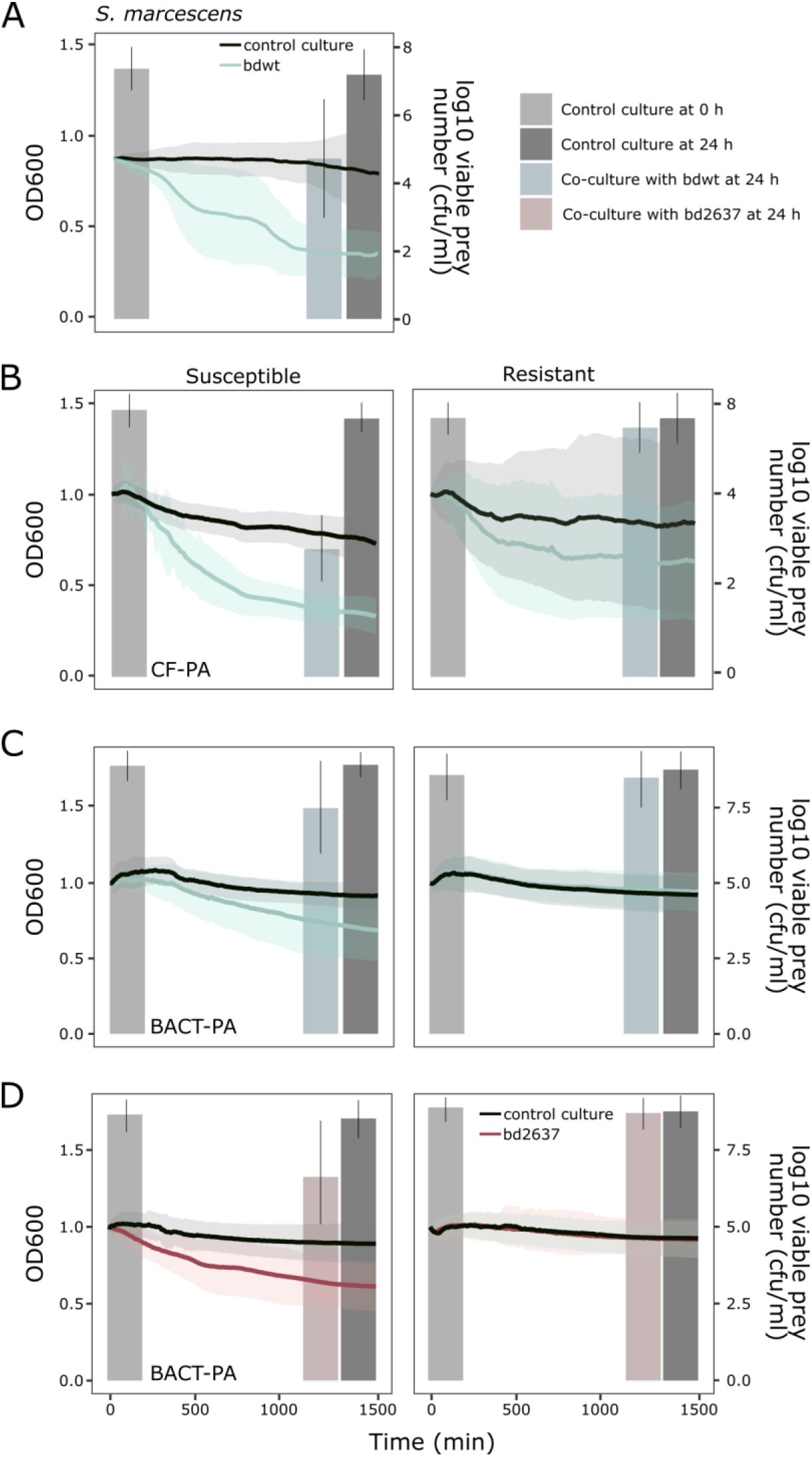
Predation susceptibility of prey collections on hepes buffer. Monitoring of OD_600_ during 24 h of predation and quantification of viable prey number. A) Predation of *B. bacteriovorus* 109J upon *S. marcescens* collection (individual predation curves in Fig S1). B) Predation of *B. bacteriovorus* 109J upon CF-PA collection (Individual predation curves in Fig S2), C) predation of *B. bacteriovorus* 109J upon BSI-PA collection (Individual predation curves in Fig S3) and D) predation of bd2637 mutant upon BSI-PA collection (Individual predation curves in Fig S6). OD_600_ was measured every 10 min and lines represent the mean of 3 biological replicates, and the shaded area indicates standard error of the mean. Bars represent the means of 3 independent viable prey quantification and error bars represent the standard error of the mean. Left and right panels represent the average of susceptible and resistant preys, respectively, of each collection.

**Fig 2.**
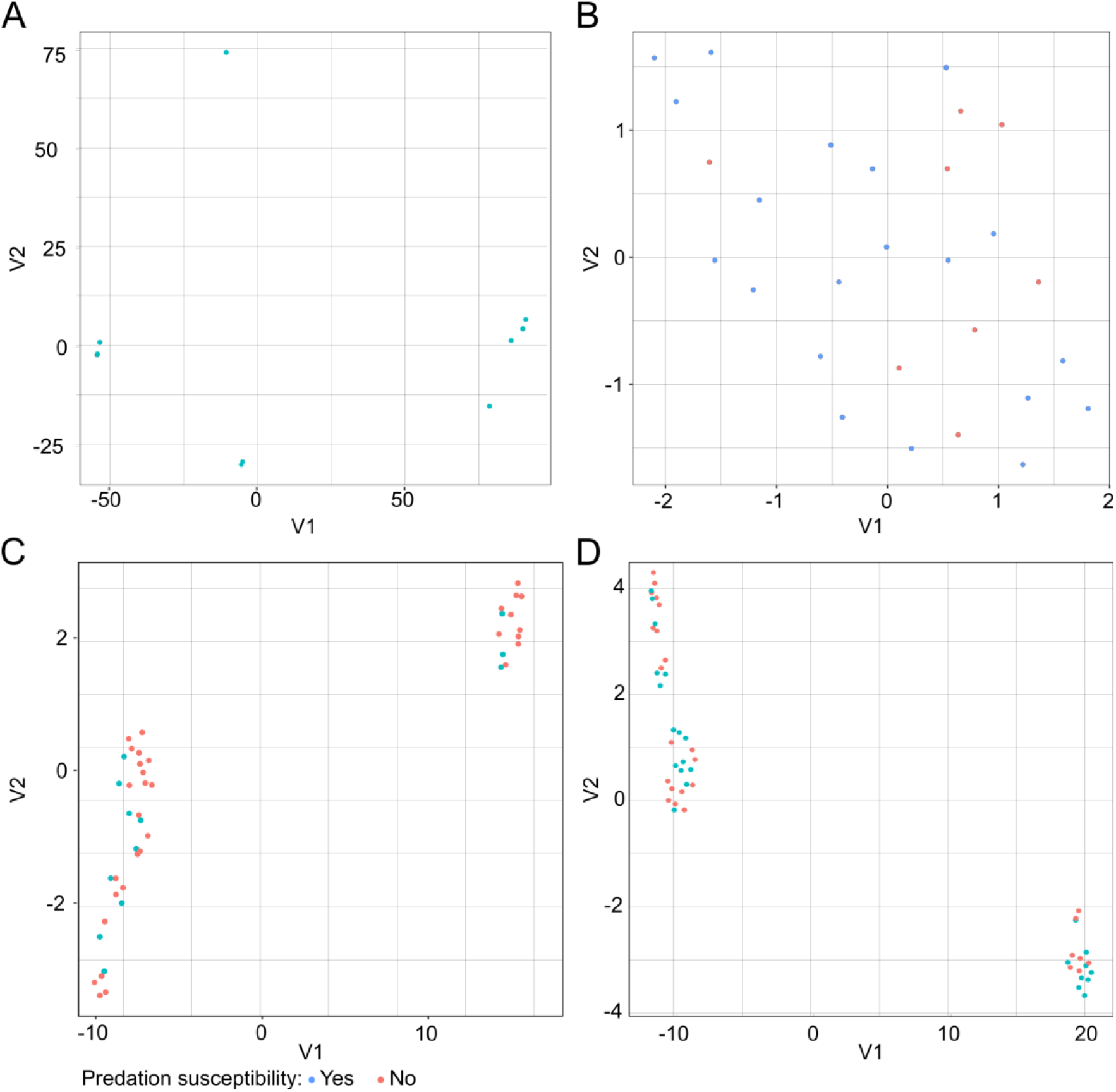
UMAP visualization of the association of predation susceptibility and antibiotic resistant profile of prey cells. A) *S. marcescens* and *B. bacteriovorus* 109J, B) CF-*P. aeruginosa* and *B. bacteriovorus* 109J, C) BACT-*P. aeruginosa* and *B. bacteriovorus* 109J; and D) BACT-*P. aeruginosa* and Bd2637 *B. bacteriovorus* data collections.

There were significant differences regarding quantitative predation (as measured by the differences in median PR) among the predation-susceptible isolates from each collection (Table S1). The median PR was 2.22 and 3.91 for *S. marcescens* and CF-PA (Mann-Whitney test *p* = 0.02), respectively, and 1.34 for BACT-PA (Mann-Whitney test *p* < 0.03). The analysis of the predation kinetics of the curves, MKR and area under the curve (AUC) did not correlate significantly with the predation rate of each isolate (Fig S5, Table S1, and Table S3).

### Influence of the predator’s predation capacity

The prey or predator determinants responsible for predation specificity have not yet been elucidated. However, the hydrolytic arsenal of *B. bacteriovorus* plays a crucial role in predation efficiency, given that it determines the success of their lifecycle (30, 41). Characterization of each predator’s specific prey spectrum is a requirement for the clinical use of predators as living antimicrobial agent. Studies have reported that different BALO lineages and predators isolated from different niches have different prey spectra (42, 43).

Interestingly, a more aggressive *B. bacteriovorus* 109J derivative has been designed and it increased by threefold the killing efficiency of the wild type preying upon *E. coli* bacterial population*s* (30). This strain was constructed by a single deletion of the gene bd2637 (coding for a PHA depolymerase) and WGS revealed that no significant chromosomal mutations were accumulated during the gene deletion process (Table S4).

We compared the predation capacity and specificity between the wild type *Bdellovibrio* and this single-gene mutant strain among the 48 BACT-PA isolates, which were the less susceptible prey collection. As expected from previous studies (30), the mutant Bd2637 strain had a higher predation frequency than the wild strain (21/48, 43.8% vs. 12/48, 25.0%); however, the differences between the PR values did not reach statistical significance (1.8 vs. 1.3, Mann-Whitney test *p* = 0.6) (Table S1, Fig 1D and S6). Interestingly, the repertoire of susceptible prey was completely different: 16 out of the 48 (33.3%) BACT-PA isolates were resistant to both predators, 15 (31.2%) were susceptible only to the mutant, 7 (14.6%) were susceptible only to the wild-type strain, and 6 (12.5%) were preyed on by both predators (Fig 3). The observed discrepancies were consistently observed in the replicates of each experiment and were not associated with any particular prey characteristic. Among the common susceptible preys (e.g., BACT1, BACT46, and BACT195), the effectiveness of each predator was strain specific. Again, the antibiotic resistance profile and the analysis of the kinetics parameters of the predation curves showed no correlation with predation susceptibility or PR (Fig 2D and S5). Thus, only the deletion of the catalytic activity derived from the bd2637 gene and the associated effects were responsible for the changes in prey susceptibility.

**Fig 3.**
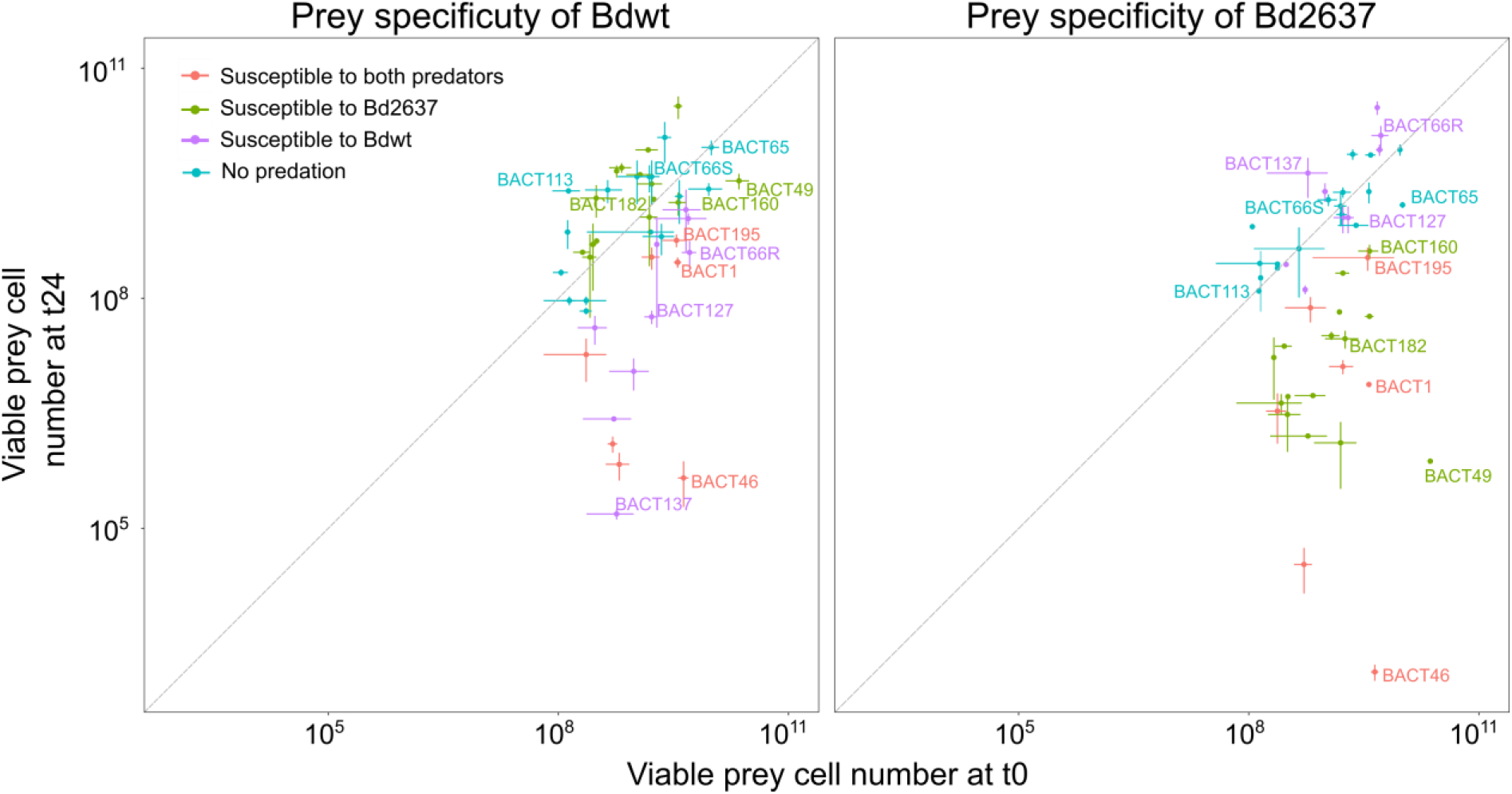
Prey specificity of predators among BACT-*P. aeruginosa* collection. Relationship between the viable prey number at the end (t24) and the beginning (t0) of the predation event. Data points below the grey line represent the susceptible preys for *B. bacteriovorus* 109J and Bd2637 *B. bacteriovorus*.

## DISCUSSION

The use of BALOs as biological control agents in environmental and medical microbiological settings (22, 44) has been suggested based on their lack of interaction with human cells (45). As occurs with antibiotics, testing the individual *in vitro* susceptibility for prey and predator pairs of strains is a requirement, mainly when predation could be substantially affected by environmental or biological conditions, as we postulate herein. Predators have been studied primarily within a free environmental context, given that the knowledge on predation susceptibility of clinical isolates is much more limited. Although human pathogens are highly diverse, we focused our research on *Pseudomonas* and *Serratia* genera due to their ubiquity in nature, their high frequency in human diseases being also carrying antibiotic resistance genes, and the availability of previously well-characterized collections including both frequent and infrequent lineages. However, the qualitative discrepancies between isolates grouped in the same ST but from different sources, indicated that the previous adaptation to these environments influences the predation process.

A single bacterial species (or ST) can be found in different habitats, as environmental and nosocomial niches (human microbiota of patients, built environments). Nevertheless, population genetic studies have revealed differing genetic evolutionary processes, in particular for those lineages highly adapted to hospital conditions, which are also known as high-risk clones, as is the case for *P. aeruginosa*, which colonizes the airways of CF-patients and are close to strains of environmental origin, as we have previously shown (46). A notable result of our study is the higher susceptibility of CF isolates to predation, without correlation with the morphological growth (mucoid vs. non-mucoid), genotype (absence of correlation with STs), or antibiotic susceptibility patterns. The presence of highly specialized predators in human microbiota cannot be ruled out, given that all available data have focused on environmental predators. Our microbial ecosystems, however, probably have the same rules of population control based on predation. Predators with a human origin could be more suitable for limiting well-adapted human pathogens (25).

In a complex and diverse ecosystem, preference for particular prey would be a dynamic feature (47). Although experiments are often conducted using individual lineages, the use of mixed populations should be a future goal to validate prey specificity in a community and the consequences on microbial population structure. This work indicates that quantitative predation, determined as relative predation rates (PR) between members of a community, could be critical to understand the dynamics of bacterial ensembles composed by different preys and predators. All *S. marcescens* isolates were susceptible to predation, as previously reported (27), and the PR was significantly higher compared with *P. aeruginosa* isolates. This finding is in line with a previous report showing a limited ability of some *B. bacteriovorus* to prey on *P. aeruginosa* (48).

The contribution of the predators’ genetic background is a pending issue. Thus far, only 8 *B. bacteriovorus* genomes have been entered into public databases, all of them from an environmental origin. To elucidate that, we used the mutant Bd2637 that was found to be a more aggressive phenotype than the wild-type *B. bacteriovorus* 109J (30). The genotype of the Bd2637 strain corresponds to a single deletion of the gen bd2637, which encodes a putative PHA depolymerase responsible for the degradation of biopolymers. The analysis of the amino acid sequence of this enzyme revealed that, apart from the characteristic esterase catalytic domain type 2 (49), the N-terminal sequence possesses a peptidase-like domain (50). This structure would provide a Bd2637 enzyme, which might act as a promiscuous enzyme with the proper catalytic architecture to act on extracellular specific components of the prey (outer membrane components, extracellular matrix, or capsid), thereby promoting specific predation. This specificity could explain why the predator is unable to complete or even begin the predatory cycle and could help identify predation resistance factors. Thus, the selection of the appropriate predator for specific prey, which needs a larger and more in-depth study on predation, would be overcome with the rational use of broad prey spectrum predators. It is worth noting that the prey range does not depend only on the prey susceptibility but also on the predator specificity, which highlights the importance of predator-prey interaction and co-evolution to overcome predation and resistance, respectively (51).

In summary, we conclude that the phenomenon of predation is defined by the particularities of both prey and predator isolates and is conditioned by environmental factors. There is a possible source dependence, and the presumption of predation cannot be inferred from different isolates of the same species, even within the same genetic lineage. To define an ecological alternative to antibiotics, the possible existence of predators within the human microbiota should be explored.

## MATERIALS AND METHODS

### Strains and growth conditions

The *B. bacteriovorus* 109J reference strain and its PHA-depolymerase mutant *B. bacteriovorus* 109J-bd2637 were used as predators in our experimental system (30). Prey (n=91) were selected from previously well-characterized collections of *S. marcescens* colonizing or infecting patients (neonates and adults) admitted to intensive care units (n=13) (31, 32). *P. aeruginosa* isolated from the airways of patients with cystic fibrosis (hereafter CF-PA, n=30) (33), and invasive isolates causing bacteremic infections (hereafter BACT-PA, n=48) (34). The inclusion criterion for the *S. marcescens* isolates, 5 of which were carbapenemase producers, was the availability of their whole-genome sequence. CF-PA and BACT-PA isolates were selected based on their genetic background discriminated by sequence type (ST), including both frequent and rare STs, as well as by their antibiotic susceptibility, including MDR isolates. Colony morphology was only considered in CF-PA isolates as mucoid (n=7) and non-mucoid isolates (n=23). Relevant data on all isolates is shown in Tables S1 and S2.

Predators were routinely grown (as described previously) by co-cultivation with *Pseudomonas putida* KT2440 in Difco nutrient broth (DNB) (0.8 g/l nutrient broth at pH 7.4) or HEPES buffer (25 mM at pH 7.8) supplemented with 2 mM CaCl_2_·2H_2_O and 3 mM MgCl_2_·3H_2_O and were further purified by filtering twice through a 0.45-μm filter. The prey strains were cultivated on Luria Broth (LB) for 16 h at 37ºC and were further diluted in HEPES buffer to an optical density at 600 nm of 1 (OD_600_ 1) for the subsequent experiments.

### Genetic background of prey and predators

Whole-genome sequencing (WGS) of both predator strains was performed on the Illumina HiSeq 4000 platform following the specification of Microbes NG (https://microbesng.com/). WGS data is available at BioProject ID: PRJNA723206 (https://www.ncbi.nlm.nih.gov/sra/PRJNA723206) and Genbank accession numbers: SRX10641169 and SRX10641170 for *B. bacteriovorus* 109J and the Bd2637 mutant, respectively. Genome comparisons were performed using *minimap2* (35) and *paftools* to align and variant calling respectively. Variant calling parameters were set to 500 bp minimum length to compute coverage and variant. Mutations were manually inspected with Artemis (36).

A tree showing the genetic relationship of 457 genomes of *S. marcescens* was constructed by combining a previously published tree (31) and 5 additional genomes of the carbapenemase-producing isolates (32). The analysis was performed by the cano-wgMLST_BacCompare web-based tool (37), and the final tree was edited using the iTOL v4.4.2 web-based tool (38). STs of the *P. aeruginosa* isolates were depicted as a minimum-spanning tree by the PHYLOViZ tool using the 7 concatenated sequences of all isolates available in May 2021 in https://pubmlst.org/organisms/pseudomonas-aeruginosa.

### Predation experiments

The predation experiments included measuring predator and prey viability and monitoring cell density (OD_600_) for 24 h. Predation co-cultures were prepared in HEPES buffer and inoculated with a *Bdellovibrio* inoculum of 10^9^ plaque-forming units/ml (PFU/ml) and a prey inoculum adjusted to an OD_600_ of 0.3. Predator and prey strain viabilities were calculated from the co-culture containing both strains. *B. bacteriovorus* strain viabilities (measured in PFU/ml) were calculated by performing serial dilutions from 10^-1^ to 10^-7^ in HEPES buffer and developing on the lawn of prey bacteria after 48–72 h of incubation at 30 °C using the double-layer method (39, 40). Briefly, 0.1 ml of the appropriate dilution was mixed with an additional 0.5 ml prey cell suspension of *P. putida* KT2440 pre-grown in LB and prepared in HEPES Buffer at pH 7.8 at OD_600_ 10, vortexed and plated on DNB solid medium. To calculate prey viable cell counts (clinical isolates), 10 μl of each dilution was plated on LB solid medium and colony-forming units (CFU/ml) were counted after 24 h of incubation at 37ºC.

Prey-predator co-cultures were performed on 96-microwell plates at a final volume of 200 μl and incubated for 24 h at 30ºC with orbital shaking in a Synergy HTX (BioTek). Two conditions were tested for each prey: the growth control well without the predator and the predation well with the mixture of prey and predator. The prey’s dynamic survival was monitored by measuring OD_600_ every 10 min, counting the viable cells at the end of the experiment by seeding on LB agar plates. Each prey was tested in at least 2–3 independent biological replicates, and the results corresponded to the mean values of all experiments.

The predation rate (PR) was calculated as the ratio of viable cells at the control well to the predation well at the end of the experiment expressed in log10 values. Positive predation is considered when this rate was >0.5. The predation kinetics encompassed the area under the curve (AUC) and the maximum killing rate (MKR). The AUC parameter was obtained using the ‘auc’ function from the ‘flux’ R package. The MKR value corresponds to the slope of the predation curve: the more negative the MKR, the more efficient the predation. The MKR was calculated as the opposite value of μmax (maximum growth rate in typical growth curves), which was obtained from the ‘growthrates’ R package. Data are represented using an R custom script and the ‘ggplot2’ package.

### UMAP clustering

UMAP was used to visualize the clustering distribution of the antibiotic resistance profile and predation susceptibility. The statistical analysis was performed in R version 3.5.0, and plotting was performed using ggplot version 2.2.1.

### Statistical analysis

Data sets were analyzed using Prism 6 software (GraphPad Software Inc.) and RStudio software v.1.2.5001. An unpaired t-test followed by Welch’s correction was performed to compare the prey’s PR. A non-parametric paired Wilcoxon test was performed to compare the MKR values in control versus predation wells, and the Kruskal-Wallis test was performed to compare PR values between collections, after assuming non-normality with the Shapiro-Wilk normality test. Lastly, the differences in predation frequency between the CF and BACT-PA collections were explored using the chi-squared test.

## ACKNOWLEDGMENTS

We appreciate the technical support of Natalia Huertas. CH is supported by Comunidad Autónoma de Madrid (PEJD-2018-POST/BMD-8016). CS is granted by “Fundación Mutua Madrileña” achieved in 2017 call by RDC (AP165902017). SSB is a recipient of a predoctoral FPU grant (FPU17/03978) from the Spanish Ministry of Universities. This work was supported by the Instituto de Salud Carlos III, PI17/00115 and PI20/00164 to RdC, REIPI (RD16/0016/0011) actions, co-financed by the European Development Regional Fund “A way to achieve Europe” (ERDF), and Vertex Pharmaceuticals.

## SUPPLEMENTARY MATERIAL

**Fig S1.**
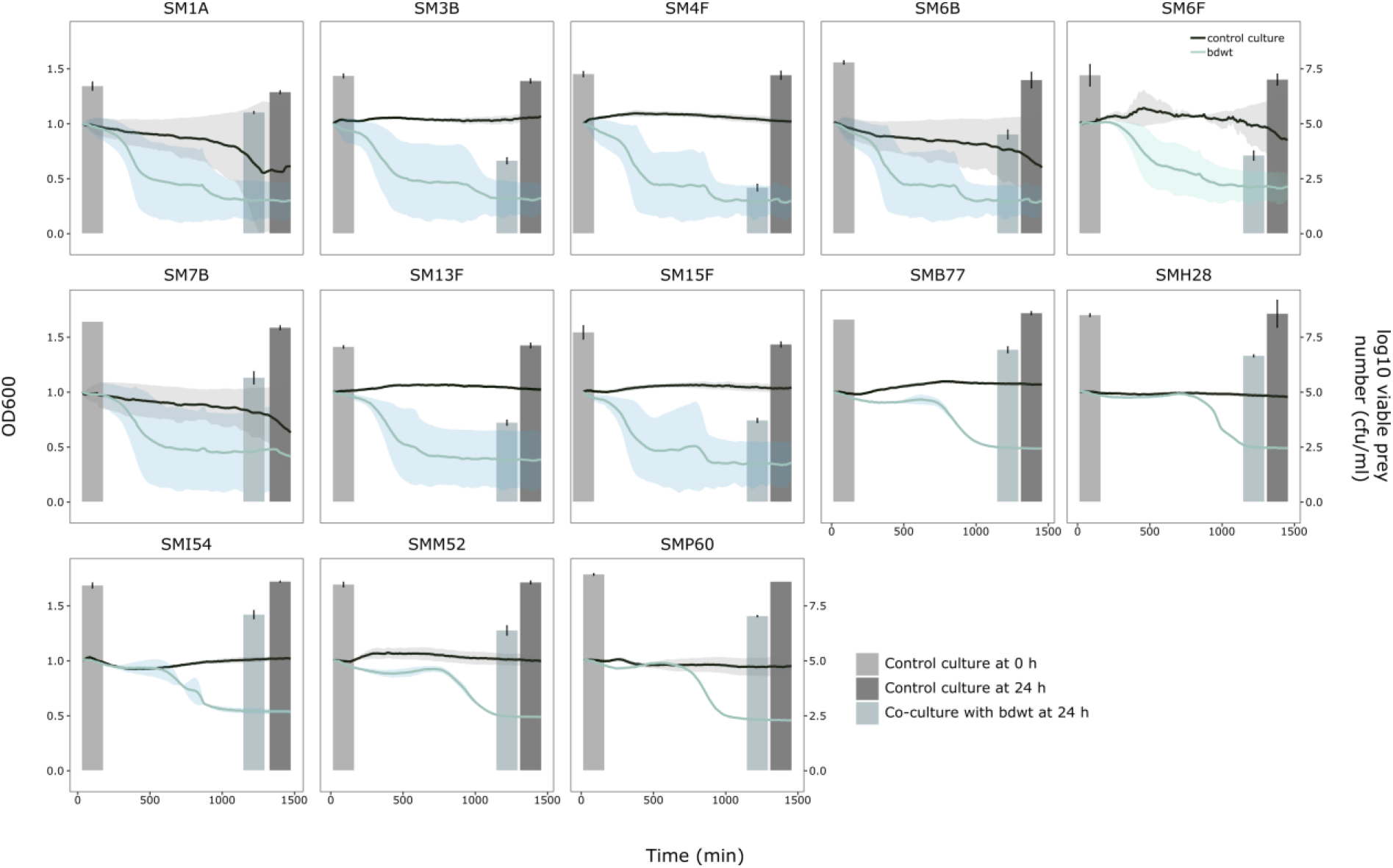
Predation susceptibility of S. marcescens prey collection by *B. bacteriovorus* 109J. OD_600_ was measured every 10 min and lines represent the mean of 3 biological replicates, and the shaded area indicates standard error of the mean. Bars represent the means of 3 independent viable prey quantification and error bars represent the standard error of the mean.

**Fig S2.**
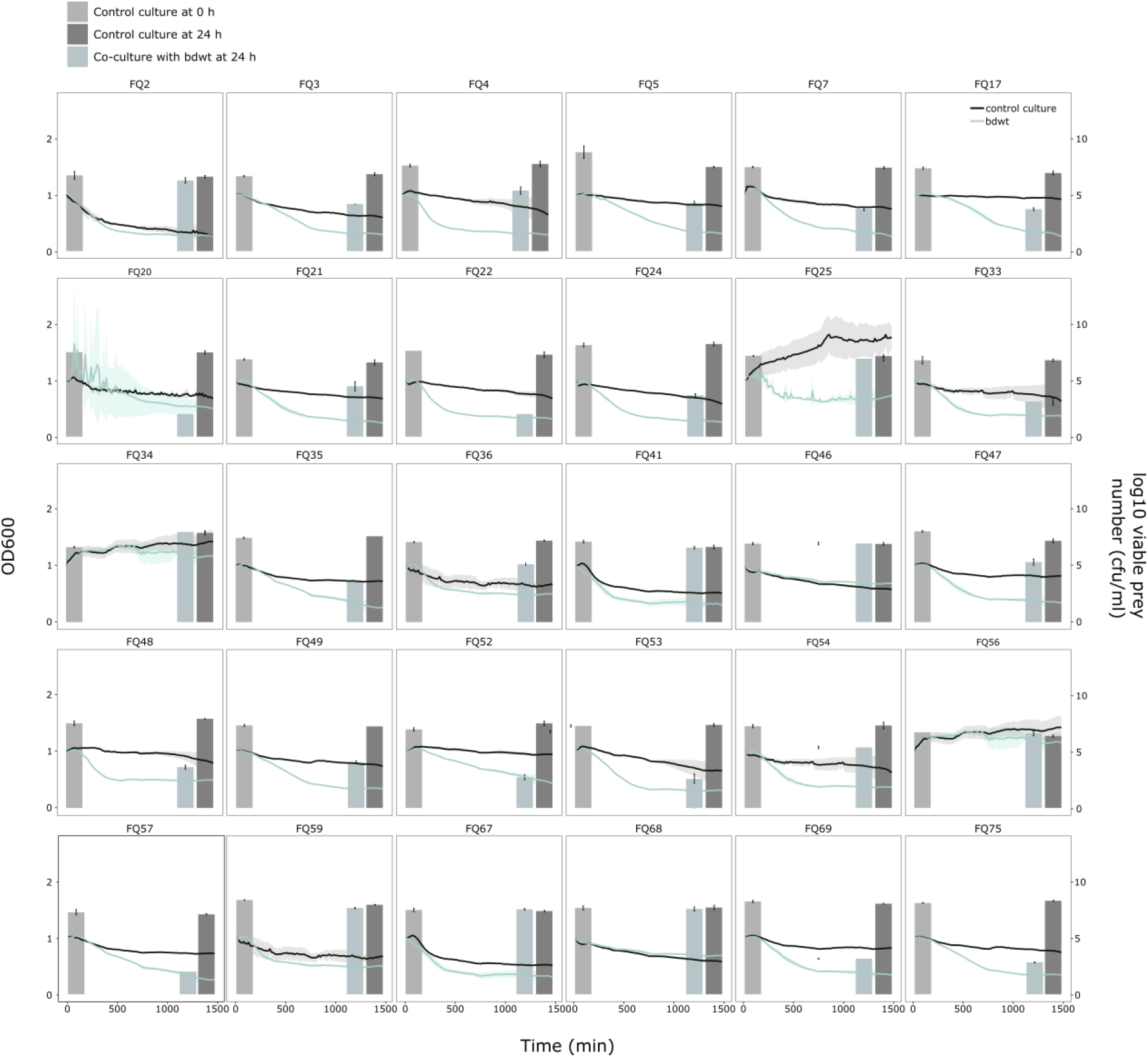
Predation susceptibility of Cf-PA prey collection by *B. bacteriovorus* 109J. OD_600_ was measured every 10 min and lines represent the mean of 3 biological replicates, and the shaded area indicates standard error of the mean. Bars represent the means of 3 independent viable prey quantification and error bars represent the standard error of the mean.

**Fig S3.**
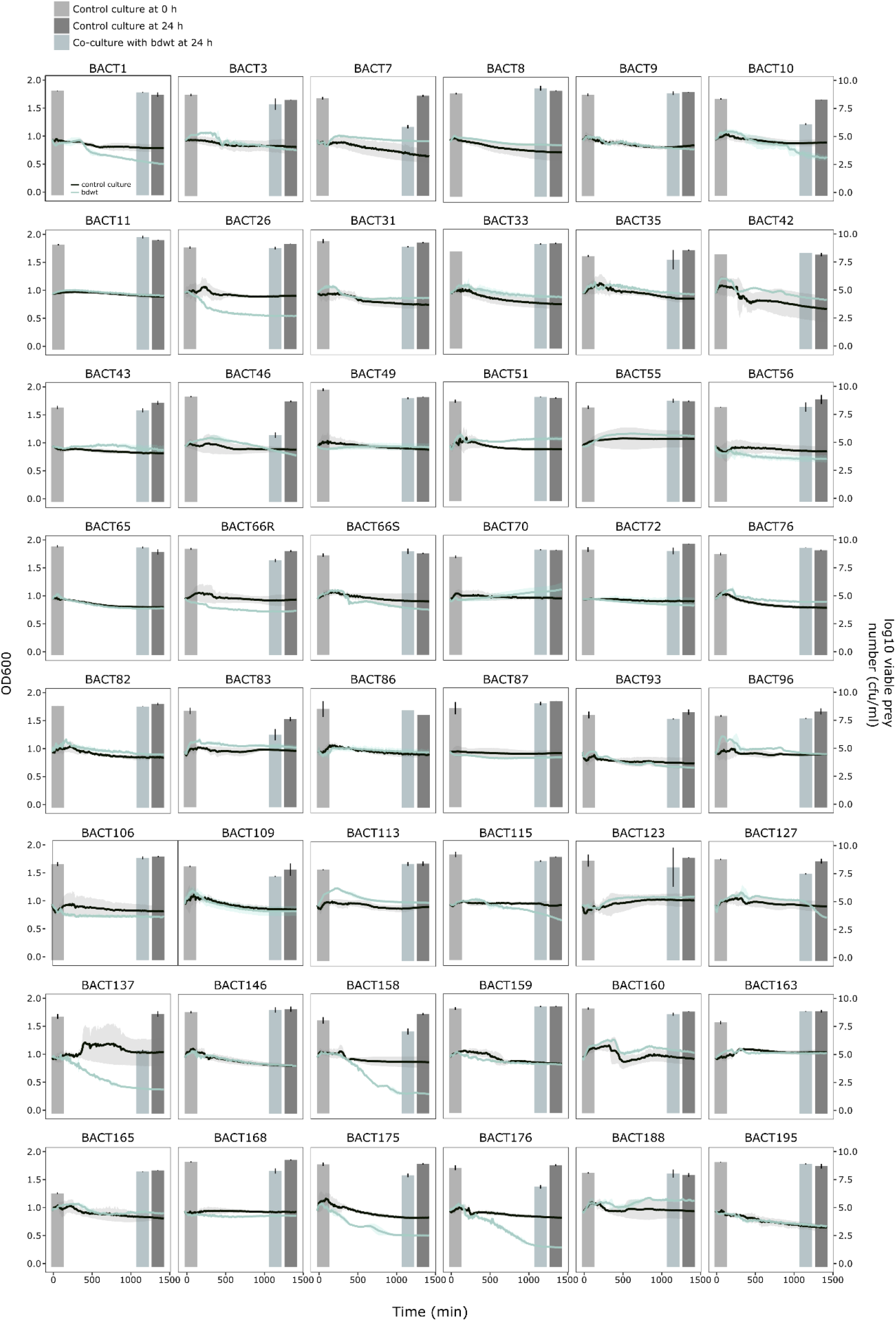
Predation susceptibility of BSI-PA prey collection by *B. bacteriovorus* 109J. OD_600_ was measured every 10 min and lines represent the mean of 3 biological replicates, and the shaded area indicates standard error of the mean. Bars represent the means of 3 independent viable prey quantification and error bars represent the standard error of the mean.

**Fig S4.**
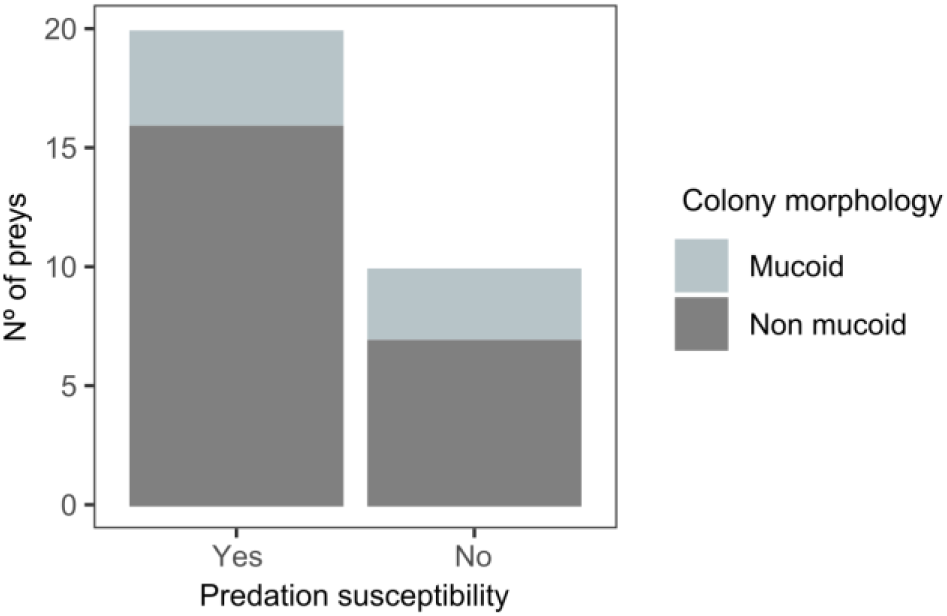
Correlation of morphotypes with predation susceptibility. Number of preys belonged to each morphotype from CF-PA collection.

**Fig S5.**
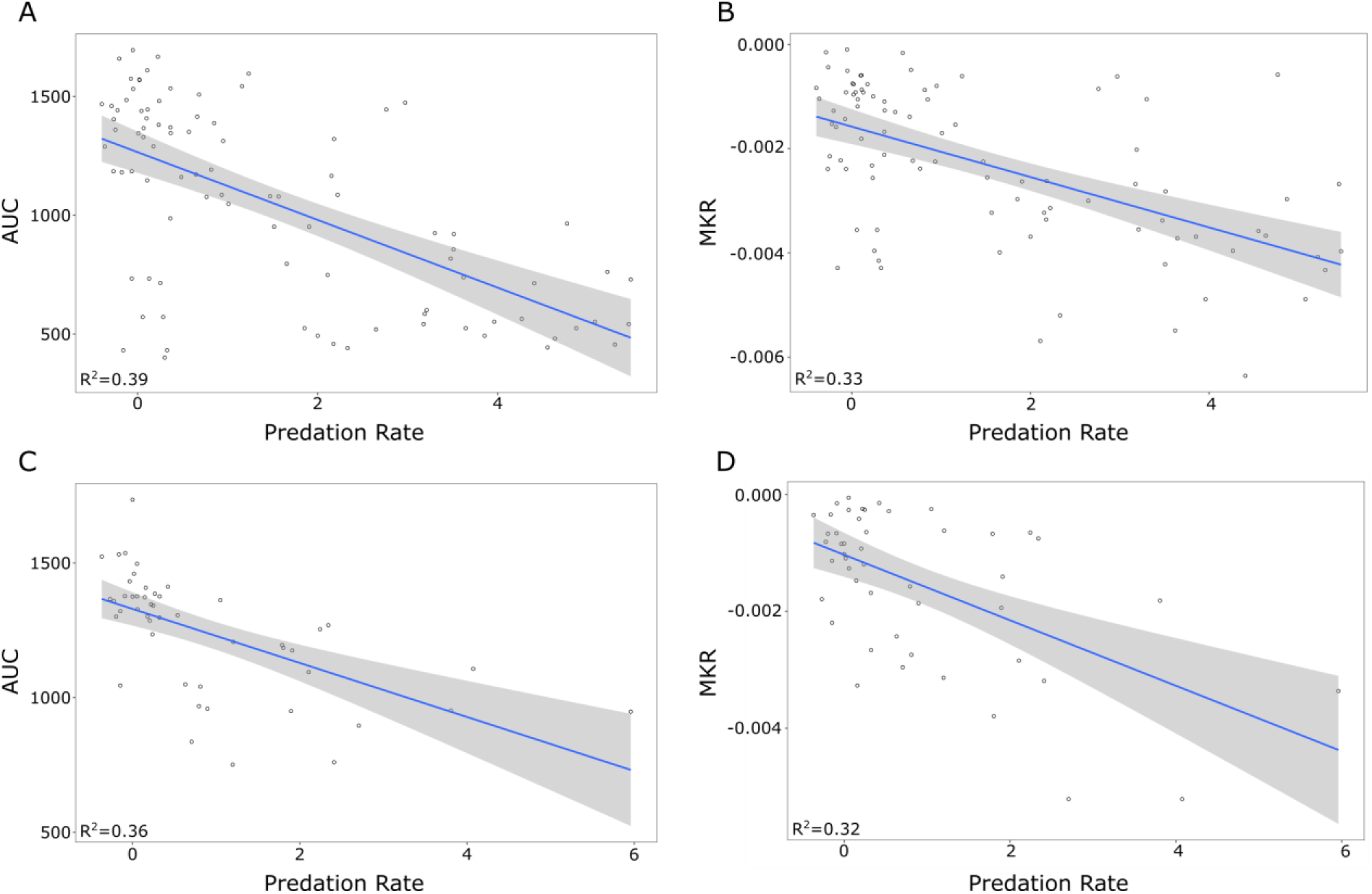
Correlation between kinetic parameter and predation rate of *B. bacteriovorus* 109J (A and B) and the mutant strain bd2637 (C and D). AUC: Area under de curve; MKR: Maximun killing rate.

**Fig S6.**
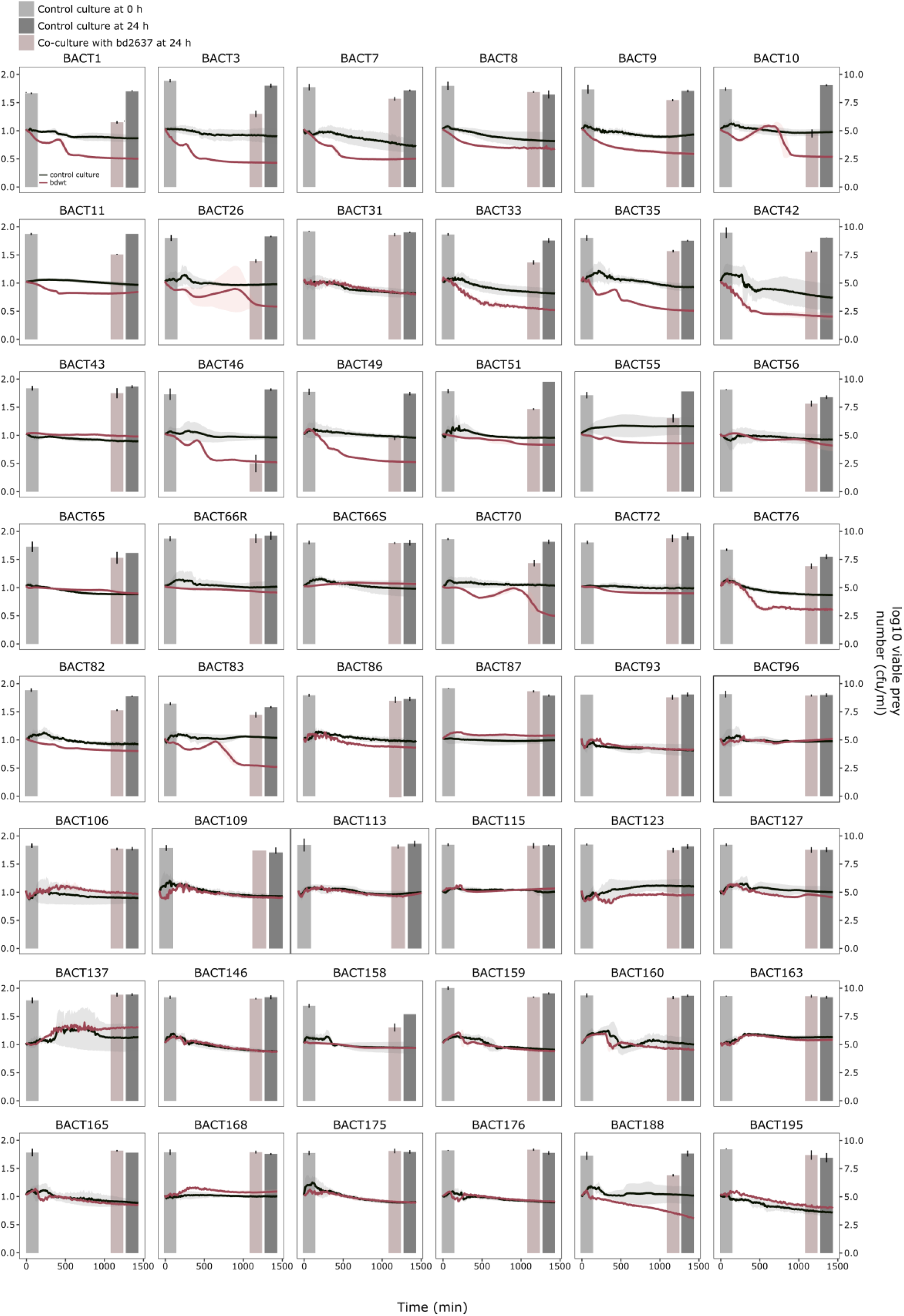
Predation susceptibility of BSI-PA prey collection by bd2637 derivative. OD_600_ was measured every 10 min and lines represent the mean of 3 biological replicates, and the shaded area indicates standard error of the mean. Bars represent the means of 3 independent viable prey quantification and error bars represent the standard error of the mean.

**Table S1. Predation susceptibility analysis of the prey clinical isolates used as prey.**

**Table S2. Antibiotic resistance profile of the clinical isolates used as prey.**

**Table S3. Kinetic parameters of predation experiments with *B. bacteriovorus* 109J and Bd2637 mutant strain**

**Table S4.**
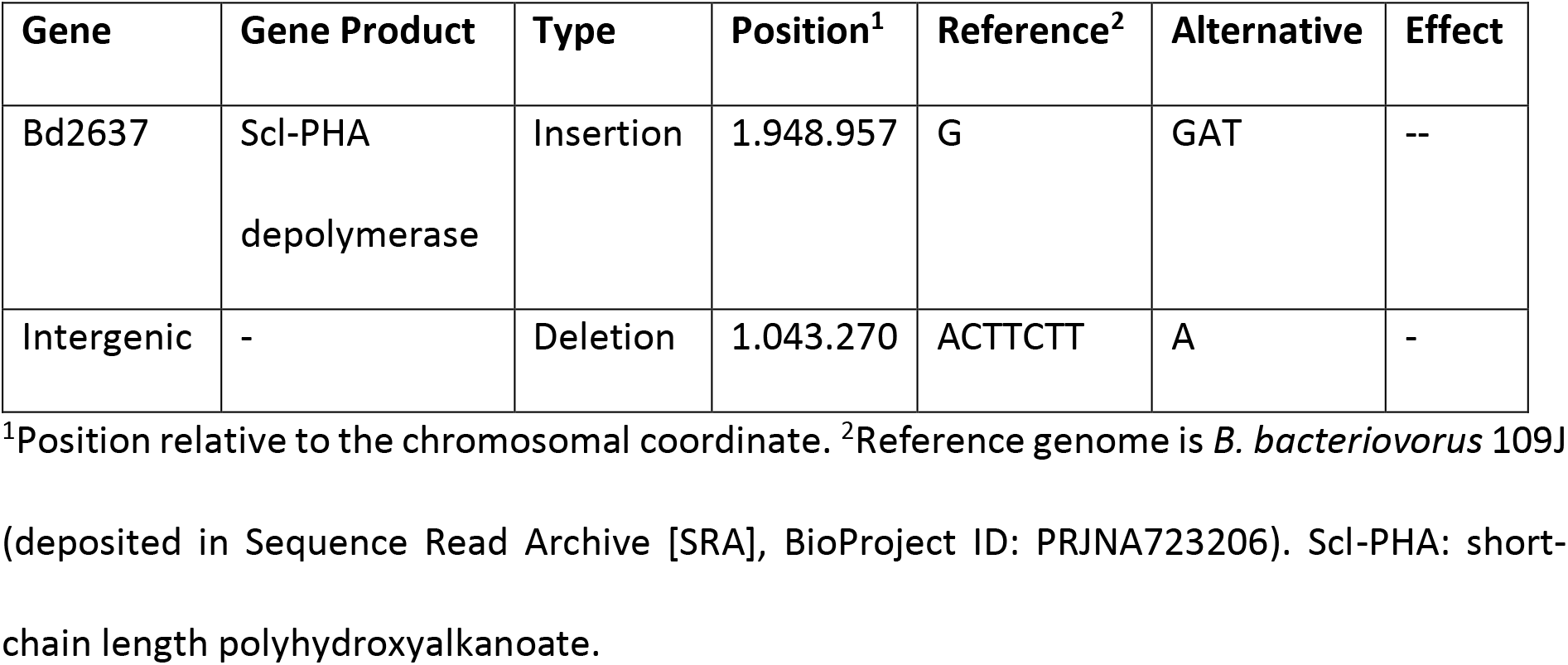
Chromosomal mutations accumulated during gene deletion.

| Gene | Gene Product | Type | Position<sup>1</sup> | Reference<sup>2</sup> | Alternative | Effect |
| --- | --- | --- | --- | --- | --- | --- |
| Bd2637 | Scl-PHA<br>depolymerase | Insertion | 1.948.957 | G | GAT | -- |
| Intergenic | - | Deletion | 1.043.270 | ACTTCTT | A | - |

<sup>1</sup>Position relative to the chromosomal coordinate. <sup>2</sup>Reference genome is *B. bacteriovorus* 109J (deposited in Sequence Read Archive [SRA], BioProject ID: PRJNA723206). Scl-PHA: short-chain length polyhydroxyalkanoate.

